# Structure of the Glutamate-Like Receptor GLR3.2 ligand-binding domain

**DOI:** 10.1101/2019.12.23.887497

**Authors:** Shanti Pal Gangwar, Marriah Green, Alexander I. Sobolevsky

## Abstract

Glutamate-like receptors (GLRs) in plants play an important role in a number of physiological processes, including wound response, stomatal aperture control, seed germination, root development, innate immune responses, pollen tube growth and morphogenesis. GLRs share amino acid sequence similarity with ionotropic glutamate receptors (iGluRs) that mediate neurotransmission in the nervous system of vertebrates. In contrast to iGluRs, however, for which numerous full-length structures are available, the structural information about the plant GLRs has been missing. Here we determine crystal structures of *Arabidopsis thaliana* GLR3.2 ligand-binding domain (LBD) in complex with glycine and methionine to 1.57 and 1.86 Å resolution, respectively. Our structures show a fold similar to iGluRs, with several secondary structure elements either missing or different. The closed clamshell conformation of GLR3.2 LBD suggests that both glycine and methionine act as agonists. The structures reveal molecular determinants of ligand binding and explain the promiscuity of GLRs’ ligand activation compared to iGluRs. Structural similarities of LBDs confirm an evolutionary relationship between GLRs and iGluRs and predict common molecular principles of their gating mechanisms that are driven by the bilobed clamshell-like LBDs.

## Introduction

Ionotropic glutamate receptors (iGluRs) are ligand-gated ion channels that mediate excitatory neurotransmission throughout the vertebrate central nervous system (CNS) (Kumar and Mayer, 2013; Traynelis et al., 2010). iGluRs assemble of 4 subunits, each containing four main domains: amino-terminal domain (ATD) implicated in receptor assembly, trafficking and regulation; ligand-binding domain (LBD or S1S2) that harbors binding sites for agonists, antagonists and, allosteric modulators; transmembrane domain (TMD) forming an ion channel; and the cytosolic carboxy-terminal domain (CTD), which is involved in receptor localization and regulation (Twomey and Sobolevsky, 2018). The neurotransmitter glutamate activates iGluR by binding to the LBD and inducing conformational changes that ultimately lead to the opening of the ion channel (Armstrong and Gouaux, 2000; Twomey and Sobolevsky, 2018). Based on pharmacology, iGluRs have been classified into four subtypes: AMPA (α-amino-3-hydroxy-5-methyl-4-isoxazolepropionic acid), NMDA (*N*-methyl-D-aspartate), kainate and δ-receptors (Traynelis et al., 2010). Interestingly, homologs of mammalian iGluRs have been identified in both vascular and non-vascular plants and are known as glutamate-like receptors (GLRs) (Lam et al., 1998).

Recent studies identified vital roles of GLRs in various physiological processes in plants, including the response to wounding, stomatal aperture control, seed germination, root development, innate immunity responses, pollen tube growth and morphogenesis (Kong et al., 2016; Kong et al., 2015; Li et al., 2013; Michard et al., 2011; Mousavi et al., 2013; Singh et al., 2016). Different number of GLRs are found in genomes of plants, including 20 for *Arabidopsis thaliana*, 2 for moss, 9 for ginkgo, 40 for pine and, 13 for tomato (Aouini et al., 2012; De Bortoli et al., 2016; Ortiz-Ramirez et al., 2017; Price et al., 2012; Wudick et al., 2018b). *Arabidopsis thaliana* GLRs (AtGLRs) are phylogenetically divided into 3 different clades (Chiu et al., 2002; Lacombe et al., 2001; Wudick et al., 2018a). AtGLR3.2, a representative of the third clade, was found localized in the plasma membrane (Vincill et al., 2013). Since overexpression of AtGLR3.2 in transgenic plants resulted in Ca^2+^ deprivation that was rescued by exogeneous Ca^2+^ application, AtGLR3.2 was proposed to function as an ion channel (Kim et al., 2001).

While GLRs govern a broad range of physiological and pathophysiological processes in plants, the molecular mechanisms underlying their function, including AtGLR3.2, remain elusive. To gain insight into how plant GLRs bind to their activating ligands, we embarked on studies of AtGLR3.2 LBD.

## Results and discussion

### Structure determination

In order to determine the LBD structure, we used *Arabidopsis thaliana* GLR3.2 (AtGLR3.2) DNA to make a crystallizing construct GLR3.2-S1S2. The boundaries of the two segments, S1 and S2, that assemble into the ligand-binding domain were determined based on the amino acid sequence alignment of AtGLR3.2 with the mammalian iGluRs (Supplementary Figure 1). Also included in the GLR3.2-S1S2 construct were 46 residues N-terminal to the beginning of S1 that have not been resolved in our crystal structures and presumably remain disordered. We expressed the GLR3.2-S1S2 construct in bacteria and purified the protein using affinity and ion-exchange chromatography (see Methods). Size-exclusion chromatography depicted the purified protein to be in monomeric form. The crystals of GLR3.2-S1S2 grew in the presence of methionine and glycine in sitting and hanging drops of vapor diffusion crystallization trays and were cryoprotected using glycerol for diffraction data collection at the synchrotron. Crystals of GLR3.2-S1S2 grown in the presence of glycine and methionine belonged to the P2_1_2_1_2_1_ space group, contained one S1S2 protomer in the asymmetric unit and diffracted to 1.57 and 1.86 Å resolution, respectively (Supplementary Table 1). We solved the GLR3.2-S1S2_Gly_ and GLR3.2-S1S2_Met_ structures by molecular replacement, initially using a homology modeled search probe (see Methods). The clarity of the resulting electron density maps were sufficient (Supplementary Figure 2) for building the structural models that included residues G47 to N286, with a 108 residue-long S1 GT-linked to a 130 residue-long S2.

The structures of approximately 57×37×35 Å^3^ in dimension have a bilobed clamshell architecture (Figure 1a-b), with the ligand-binding site between the upper D1 lobe and the lower D2 lobe, similar to the iGluR LBDs (Gouaux, 2004). The GLR3.2-S1S2_Gly_ and GLR3.2-S1S2_Met_ structures superpose very well with the root mean square deviation (RMSD) of 0.275 Å for Cα atoms. Despite the GLR3.2-S1S2_Gly_ and GLR3.2-S1S2_Met_ backbone similarity, several residues show different side chain orientations, including Asp68, Lys132, Glu145, Arg159 and Arg281, with the greatest difference in side-chain rotamers observed for Arg281. For the ligand-binding pocket, however, side-chain orientations are very similar between GLR3.2-S1S2_Gly_ and GLR3.2-S1S2_Met_.

**Figure 1.**
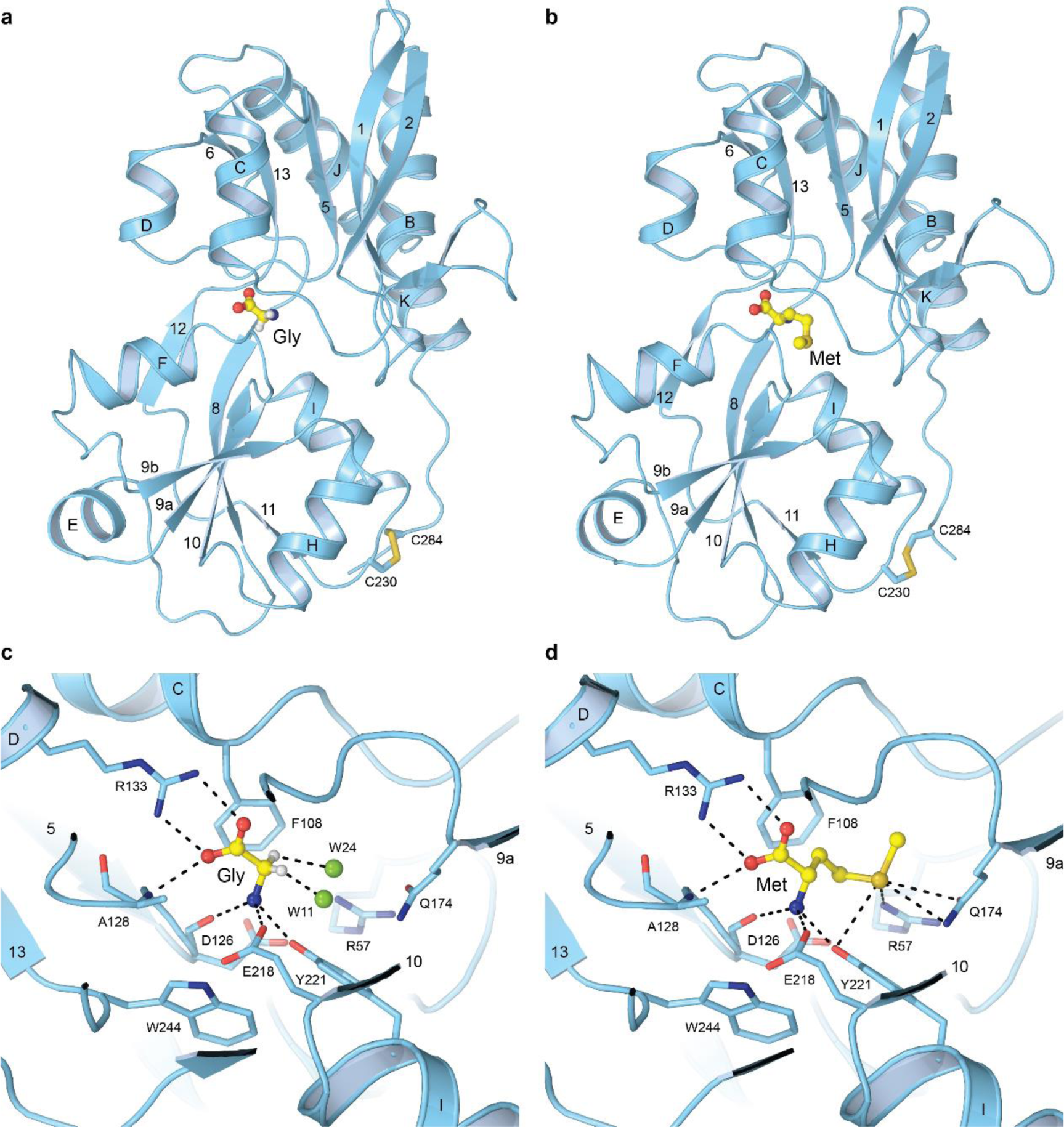
GLR3.2 ligand-binding domain structure. **a-b**, Structures of isolated GLR3.2 LBD (S1S2) in complex with glycine (**a**) and methionine (**b**). The ligands are in ball-and-stick representation. Highly conserved cysteines C230 and C284 connected by disulfide bond are shown in sticks. **c-d**, Close-up views of the ligand-binding pocket with bound glycine (**c**) and methionine (**d**). Residues involved in ligand binding are shown in sticks. Characteristic interactions between the ligands and the binding pocket residues are indicated by dashed lines.

**Figure 2.**
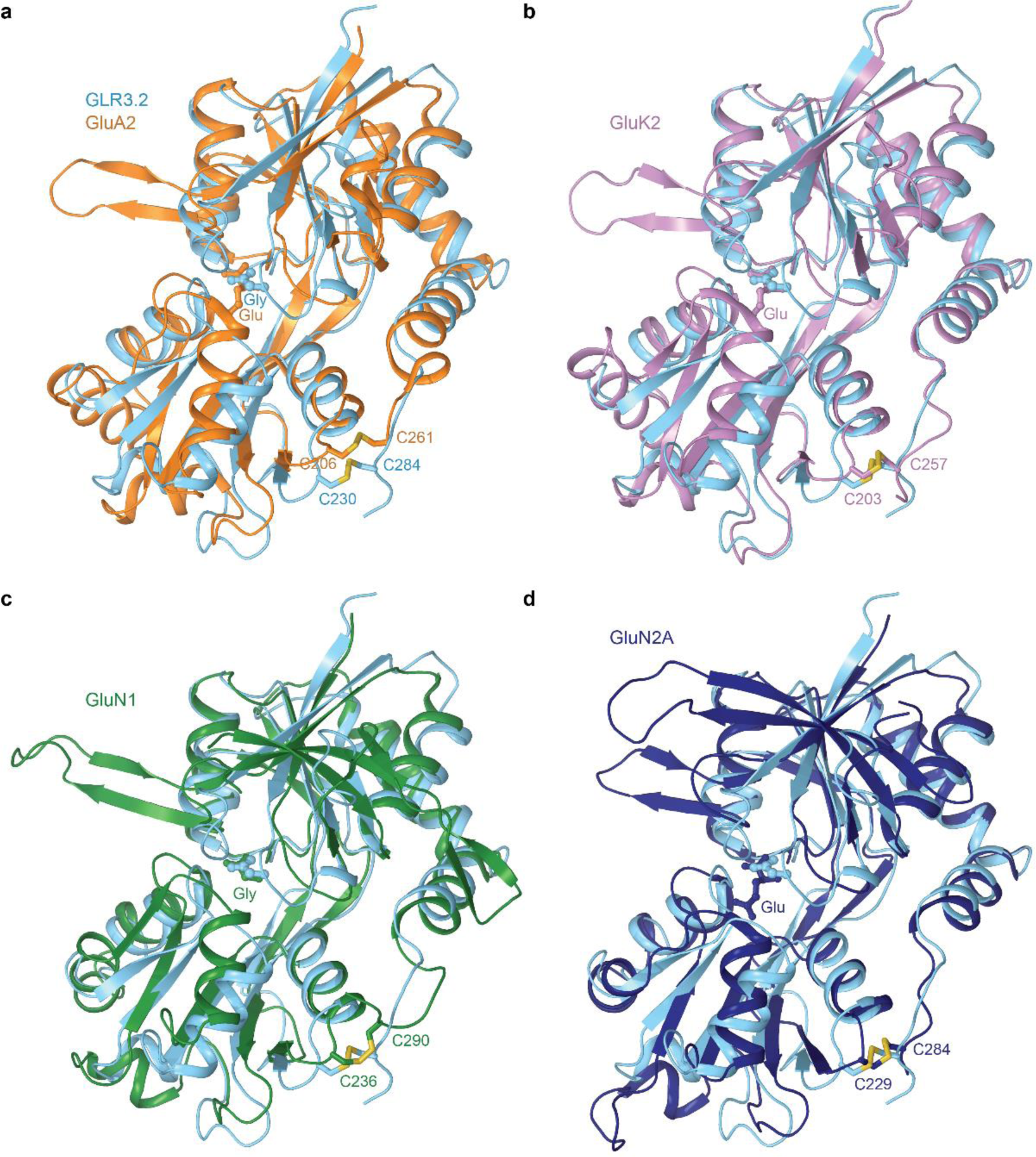
Comparison of GLR3.2 and iGluR LBDs. **a-d**, Structural superpositions of isolated LBDs from GLR3.2 (cyan) in complex with glycine and (**a**) rat GluA2 (PDB ID: 1FTJ, orange) in complex with glutamate, (**b**) rat GluK2 (PDB ID: 1S50, purple) in complex with glutamate, (**c**) rat GluN1 (PDB ID: 1PB7, green) in complex with glycine and (**d**) rat GluN2A (PDB ID: 2A5S, blue) in complex with glutamate. The ligands are in ball-and-stick representation. Highly conserved cysteines connected by disulfide bonds are shown in sticks.

### Ligand binding

The ligand-binding pocket of GLR3.2 LBD also resembles the ligand-binding pocket of iGluR LBDs (Figure 1c-d), with the key interactions and binding residues conserved (Supplementary Figure 1). The total of seven residues, Arg57, Asp126, Ala128, Arg133, Gln174, Glu218 and Tyr221, contribute primarily to the LBD-ligand interactions. For both glycine and methionine, the guanidinium group of Arg133 and the backbone amine of Ala128 coordinate the carboxyl group of the ligand, while the backbone carbonyl oxygen of D126, the carboxyl group of Glu218 and hydroxyl group of Tyr221 coordinate the amino group of the ligand. The thioether group of methionine is additionally coordinated by the hydroxyl group of Tyr221, guanidinium group of Arg57 and the amide group of Gln174. These interactions are specific to methionine and are missing in case of glycine, which lacks the bulky side chain. Instead, two water molecules occupy the space that in case of methionine is occupied by the thioether group. Water molecules are hydrogen bonded to glycine’s Cα, thus stabilizing binding of this ligand to LBD.

Overall, the ligand-binding pocket of GLR3.2 is shaped to universally bind differently sized amino acids (for example, glycine versus methionine) by exploiting the same interactions for binding the conserved amino acid core and adjusting the fit of the side chain into the corresponding biding pocket cavity with water. This explains a diverse range of ligand specificity previously observed for GLRs, with at least 12 of the 20 proteinogenic amino acids and D-Serine serving as agonists for the most studied AtGLR1.2, AtGLR1.4, AtGLR3.3, AtGLR3.4, and AtGLR3.5(Forde and Roberts, 2014; Kong et al., 2016; Michard et al., 2011; Tapken et al., 2013; Vincill et al., 2012; Vincill et al., 2013; Wudick et al., 2018a). Such promiscuity has never been observed in iGluRs. The binding pocket and the mode of ligand binding, however, might be somewhat different between GLRs. For example, Trp, Phe and Tyr can serve as agonists of AtGLR1.4 but not AtGLR3.3 or AtGLR3.4 (Tapken et al., 2013; Vincill et al., 2012; Vincill et al., 2013) suggesting that the ligand-binding pocket in AtGLR1.4 is likely larger and can accommodate bulky hydrophobic side chains.

### Comparison of GLR3.2 and iGluR LBD structures

The GLRs share poor sequence identity to iGluRs, nevertheless, the structural fold of their LBDs is highly conserved. However, there are some major structural differences between the GLR and iGluR LBDs. For structural comparison, we superimposed the GLR3.2 LBD with the previously solved agonist-bound LBD structures of GluA1 (Armstrong and Gouaux, 2000), GluK2 (Mayer, 2005), GluN1 (Furukawa and Gouaux, 2003) and GluN2A (Furukawa et al., 2005). The overall LBD fold is quite similar between GLR3.2 and iGluRs. The RMSD value for all Cα atoms is 1.9, 1.8, 1.5 and 4.5 Å for GluA1, GluN1, GluK2 and GluN2, respectively. In all iGluRs, the highly conserved and structurally important disulfide bond formed between the regions C-terminal to helices I and K is also present in the GLR3.2 LBD and connects Cys230 to Cys284. Compared to iGluR LBDs, however, the GLR3.2 LBD is missing the β-hairpin loop between β2 and helix C. Neither GLR3.2 LBD has the helix G. Instead, this region adopts a short β strand that we named 9b. The loop between β1 and helix B is shorter in GLR in comparison to GluN1 (1PB7), GluK2 (1S50) and GluN2A (2A5S). This region harbors the ligand-binding Arg57 residue. In addition, NMDA receptor LBDs also have a loop between β1 and helix B, forming a large hairpin, which is missing in GLR3.2 as well as AMPA and kainate receptor LBDs.

Critically, the extent of the LBD clamshell closure around the ligand is very similar in LBDs of GLR3.2 and all agonist-bound iGluRs. This suggests that both glycine and methionine play a role of agonists or partial agonists of GLR3.2, which may explain previous reports of GLRs (GLR3.1/3.5) as L-Met-activated Ca^2+^ channels responsible for maintaining cytosolic Ca^2+^ (Kong et al., 2016). Accordingly, the LBD captured in our GLR3.2-S1S2_Gly_ and GLR3.2-S1S2_Met_ structures might act similar to the LBD in iGluRs by opening their clamshell in the absence of a ligand and closing it in its presence, where the clamshell closure provides a driving force to gate an ion channel (Armstrong and Gouaux, 2000; Twomey and Sobolevsky, 2018). To test this hypothesis, one would need to capture the full-length structure of GLR. The predicted similarity in the LBD clamshell architecture, ligand binding and gating mechanism also suggests that plant GLRs and iGluRs originated from a common ancestor to function in different kingdoms of life but utilizing similar molecular mechanisms.

## Acknowledgments

We thank Dr. Surajit Banerjee for assistance with the data collection, Dr. Jesse Yoder for help with the molecular replacement, and Dr. Appu K. Singh for advice in the crystallographic data processing. We thank Drs. Maria Yelshanskaya and Kirill Nadezhdin for comments on the manuscript and for helpful discussions. A.I.S. is supported by the NIH (R01 CA206573, R01 NS083660, R01 NS107253), NSF (1818213) and the Irma T. Hirschl Career Scientist Award. Data was collected at the beamline 24-ID-C of the Advanced Photon Source. 24-ID-C is one of the Northeastern Collaborative Access Team beamlines, which are funded by the National Institute of General Medical Sciences from the National Institutes of Health (P30 GM124165). The Pilatus 6M detector on the 24-ID-C beamline is funded by an NIH-ORIP HEI grant (S10 RR029205).

## Author Contributions

A.I.S. supervised the project. S.P.G. and M.G. made constructs and prepared protein samples. S.P.G. and A.I.S. carried out crystallographic data collection, processing and built molecular models. S.P.G., M.G., and A.I.S. wrote the manuscript.

## Methods

### Construct

DNA for *Arabidopsis thaliana* GLR3.2 was a gift of Prof. José A. Feijó (University of Maryland). The boundaries of the GLR3.2 ligand-binding domain (S1S2) were determined based on the sequence alignment with GluA2 (Armstrong et al., 1998; Sobolevsky et al., 2009). The DNA encoding AtGLR3.2 residues Ser420-V572 (S1) and P682-N811 (S2) was amplified using gene-specific primers and subcloned into the pET22b vector (Novagen) between NcoI and XhoI sites with a GT linker between S1 and S2 (Armstrong and Gouaux, 2000). For purifications purposes, 8xHis affinity tag followed by a thrombin cleavage site (LVPRG) were introduced N-terminally.

### Protein expression and purification

The construct pET22b carrying GLR3.2-S1S2 was transformed into *Escherichia coli* Origami B (DE3) cells and grown in LB media supplemented with 100 µg/ml ampicillin, 15 µg/ml kanamycin and 12.5 µg/ml tetracycline. The freshly inoculated culture was grown at 37°C until OD_600_ reached the value of 1.0-1.2. Then cells were cooled down to 20°C, induced with 250 µM IPTG and incubated in the orbital shaker for another 20 hours at 20°C. Cells were harvested by centrifugation at 5000 rpm for 15 min at 4°C and the cell pellet was washed with the buffer containing 20 mM Tris pH 8.0 and 150 mM NaCl. For protein extraction, cells were resuspended in lysis buffer consisting of 20 mM Tris pH 8.0, 200 mM NaCl, 1 mM glutamate, 5 mM methionine, 1 mM βME, 1 mM PMSF, 100 µg/ml lysozyme, 5 mM MgSO_4_ and DNAse. All purification steps were carried out in buffers supplemented with 1 mM glutamate and 5 mM methionine. The cells were disrupted by sonication and centrifuged at 40,000 rpm in the Ti45 rotor for 1 hour at 4°C. The supernatant was mixed with His60 Ni superflow resin (Takara) and rotated for 2 hours at 4°C. The protein-bound resin was washed with the buffer containing 15 mM imidazole and the protein was eluted in 20 mM Tris pH 8.0, 150 mM NaCl, 1 mM glutamate, 5 mM methionine, 1 mM βME, and 200 mM Imidazole. The protein was dialyzed overnight in the buffer containing 20 mM Tris pH 8.0, 75 mM NaCl, 1 mM glutamate, 5 mM methionine, 1 mM BME, and 4% glycerol. After thrombin digest (1:500 w/w) at 22°C for 1-hour, the protein was further purified using ion-exchange Hi-Trap Q HP-(GE Healthcare). The protein quality was assessed by SDS-PAGE and analytical size-exclusion chromatography using the Superpose 10/300 column (GE Healthcare).

### Crystallization and structure determination

Crystallization screening was performed with GLR3.2-S1S2 protein at a concentration of ~7 mg/ml using Mosquito robot (TTP Labtech) and sitting drop vapor diffusion in 96-well crystallization plates. Small needle-shaped crystals, which appeared after two weeks of incubating crystallization trays at 4°C and 20°C, were further optimized using the hanging drop method and 24-well crystallization plates. The best-diffracting long needle-shaped crystals of methionine-bound GLR3.2-S1S2 grew at 20°C in 0.1 M MES pH 6.5, 18% PEG MME 2K and 0.1 M ammonium sulfate. Crystals of glycine-bound GLR3.2-S1S2 grew in a similar condition but in the presence of 0.3 µl of 1M glycine that supplemented the 4µl crystallization drop as an additive. The best-diffracting needle-shaped crystals of glycine-bound GLR3.2-S1S2 grew at 4°C in 22 % PEG 4K, 0.1 M ammonium acetate, and 0.1 M sodium acetate pH 4.6. All crystals were cryoprotected using 25% glycerol and flash-frozen in liquid nitrogen for data collection. Crystal diffraction data were collected at the beamline 24-ID-C of the Advanced Photon Source and processed using XDS (Kabsch, 2010) and Aimless as a part of the CCP4 suite (Winn et al., 2011).

The structure of methionine-bound GLR3.2-S1S2 was solved by molecular replacement using Phaser (McCoy, 2007) and a search probe generated by SWISS-MODEL homology modeling (Waterhouse et al., 2018) from the ligand-binding domain of NMDA receptor (PDB ID: 6MMS) (Jalali-Yazdi et al., 2018). The initial partial solution was used again as a search probe for subsequent rounds of molecular replacement, which ultimately resulted in a complete GLR3.2-S1S2 model. The model was refined by alternating cycles of building in COOT (Emsley and Cowtan, 2004) and automatic refinement in Phenix (Adams et al., 2010). The structure of glycine-bound GLR3.2-S1S2 was solved by molecular replacement using the methionine-bound GLR3.2-S1S2 structure as a search probe. Water molecules were added in Coot and Phenix refine. All structural figures were prepared in PyMol (DeLano, 2002).

### Data Availability

Model coordinates have been deposited to the Protein Data Bank (PDB) under accession numbers XXXX for GLR3.2-S1S2 structure in complex with glycine and YYYY in complex with methionine. All other data are available from the corresponding author upon request.

---

**Supplementary Figure 1.**
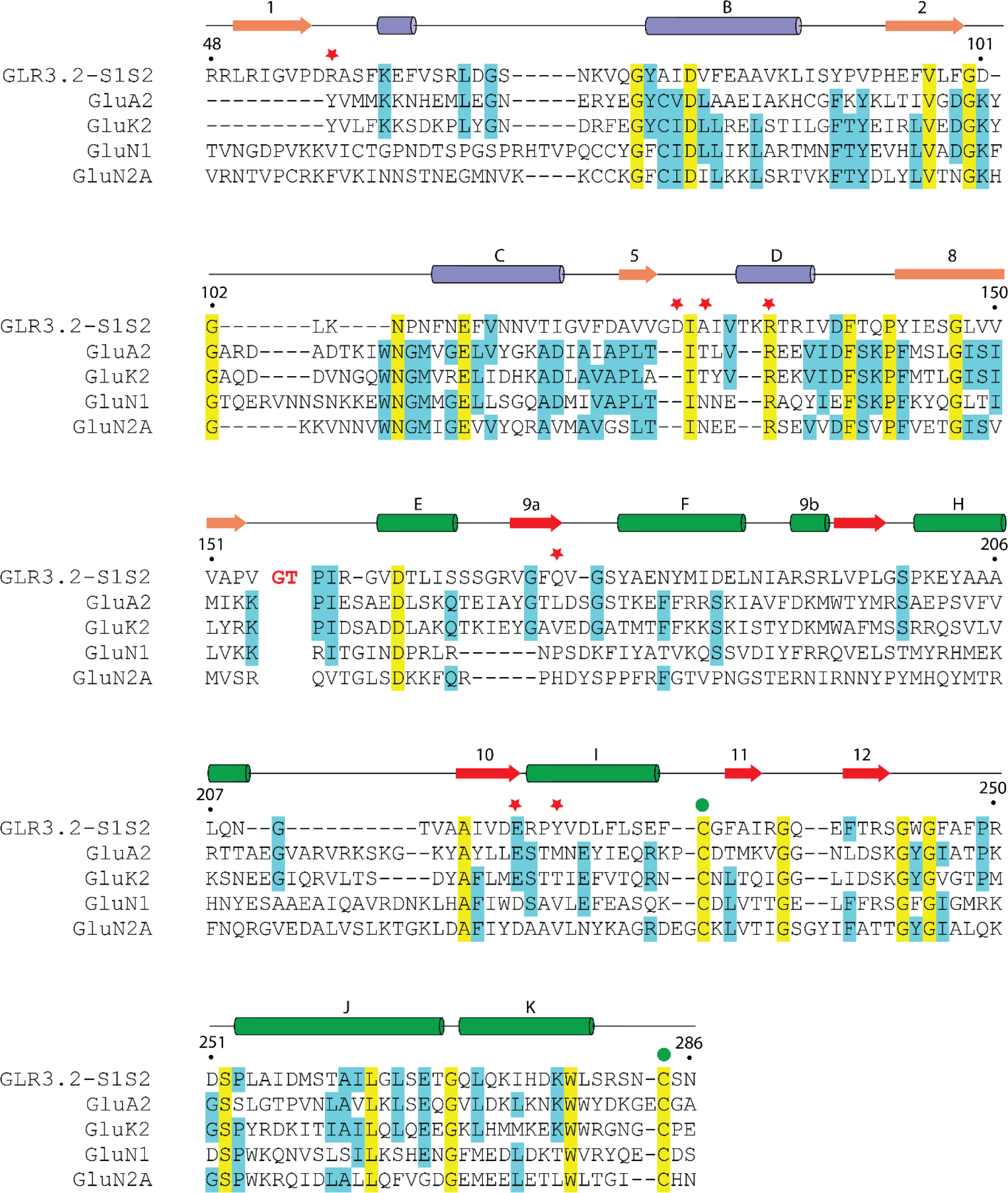
Amino acid sequence alignment. Shown are amino acid sequences for the GLR3.2-S1S2 construct and ligand-binding domains of AMPA subtype rat GluA2 (NP_058957), kainate subtype rat GluK2 (P42260.2), and NMDA subtype rat GluN1 (EDL93606.1) and GluN2A (NP_036705.3) subunits. Numbering is for the mature protein. Secondary structure elements for GLR3.2-S1S2 are shown as cylinders (α-helices), arrows (β-strands) and lines (loops) colored according to domains S1 (orange and purple) and S2 (red and green). The names of α-helices and β-strands (capital letters and numbers, respectively) are kept the same as in structures of isolated LBD (Armstrong et al., 1998). Completely conserved residues are highlighted in yellow and mostly conserved residues are highlighted in blue. Green circles indicate cysteines connected by disulfide bonds. Red stars indicate residues involved in ligand binding.

**Supplementary Figure 2.**
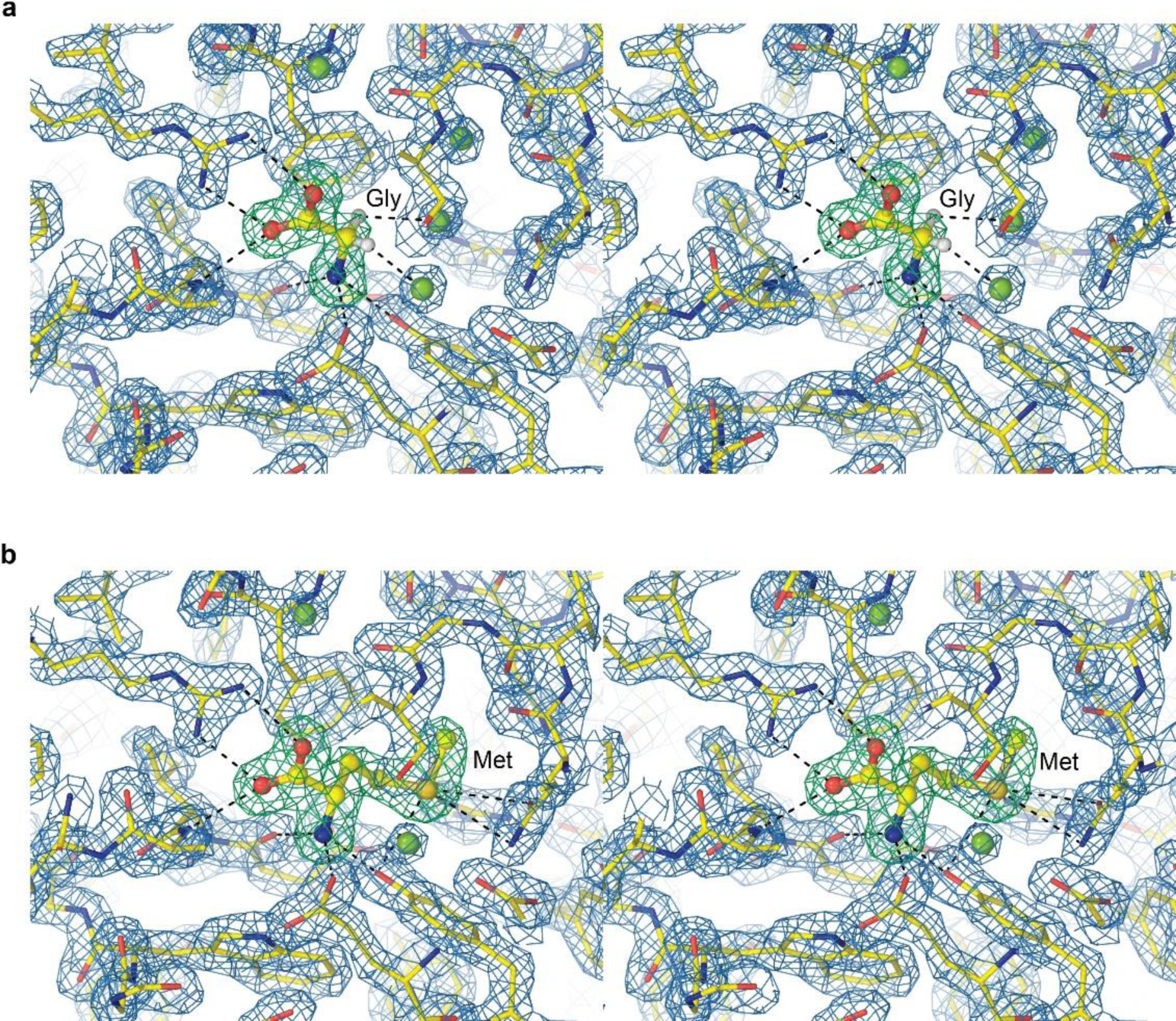
GLR3.2-LBD electron density. **a-b**, Close-up stereo view of the ligand-binding domain in isolated GLR3.2 LBD (S1S2) in complex with glycine (**a**) and methionine (**b**). Mesh shows 2Fo-Fc electron density map contoured at 2 σ (blue) and Fo-Fc map contoured at 4 σ (green) when ligands were not present in the model.

**Supplementary Table 1.** Crystallographic statistics.

|  | **GLR3.2-S1S2<sub>Gly</sub>** | **GLR3.2-S1S2<sub>Met</sub>** |
| --- | --- | --- |
| Beamline | NE-CAT 24-ID-C | NE-CAT 24-ID-C |
| Wavelength (Å) | 0.97910 | 0.97910 |
| Space group | P2<sub>1</sub>2<sub>1</sub>2<sub>1</sub> | P2<sub>1</sub>2<sub>1</sub>2<sub>1</sub> |
| Cell parameters (a, b, c, Å) | 47.39, 64.37, 75.93 | 48.40, 65.74, 73.19 |
| Cell parameters ($\alpha$, $\beta$, $\gamma$, °) | 90, 90, 90 | 90, 90, 90 |
| Resolution (Å) | 40.2-1.57 (1.62-1.57) | 48.91-1.86 (1.90-1.86) |
| Number of Monomers in AU | 1 | 1 |
| Observations | 211680 (17139) | 115986 (3554) |
| Unique reflections | 33057 (3206) | 19714 (1516) |
| R<sub>merge</sub> | 0.06 (0.78) | 0.09 (0.15) |
| R<sub>mease</sub> | 0.07 (0.86) | 0.11 (0.20) |
| R<sub>pim</sub> | 0.02 (0.36) | 0.04 (0.11) |
| Mean (I)/sigma (I) | 17.9 (2.1) | 19.3 (5.6) |
| Completeness (%) | 99.6 (98.3) | 96.9 (76.5) |
| Multiplicity | 6.4 (5.3) | 5.9 (2.3) |
| Wilson B-factors (Å<sup>2</sup>) | 18.9 | 17.9 |
| **Refinement** |  |  |
| Reflections used in refinement | 33024 (3206) | 19644 (1515) |
| R<sub>work</sub> | 0.155 (0.233) | 0.148 (0.176) |
| R<sub>free</sub> | 0.186 (0.264) | 0.182 (0.214) |
| Number of protein atoms | 1861 | 1833 |
| Number of ligands | 1 | 1 |
| No of ions | 0 | 4 |
| Average B factor | 23.7 | 24.8 |
| Number of water molecules | 414 | 341 |
| RMSD bond lengths (Å) | 0.01 | 0.01 |
| RMSD angles (°) | 1.03 | 0.81 |
| **Ramachandran plot** |  |  |
| Preferred regions (%) | 98.32 | 99.57 |
| Allowed regions (%) | 0.84 | 0.00 |
| Outliers (%) | 0.84 | 0.43 |

Values in parentheses are for the highest-resolution shell.

